# Inhaled black carbon induces depressive-like behavior and enhances stress-related blood–brain molecular vulnerability in mice

**DOI:** 10.64898/2026.08.24.745657

**Authors:** Jinhee Bae, Jiwon Lee, Seongeun Song, Kwangil Jeong, Nazarii Frankiv, Chaeran Park, Chae Yeon Hwang, Yun Kyung Kim, Byung-Yong Yu, Heh-In Im

**Author notes:** Correspondence: Heh-In Im, Ph.D., Brain Disease Research Center, Brain Science Institute, Korea Institute of Science and Technology, 5 Hwarang-ro 14-gil, Seongbuk-gu, Seoul 02792, Republic of Korea.

## Abstract

Black carbon (BC), a combustion-derived component of fine particulate matter, has been linked to depressive symptoms, but controlled experimental evidence remains limited. We established a controlled BC inhalation model combined with chronic restraint stress (CRS) to determine whether inhaled BC alone induces depressive-like behavior and whether concurrent stress enhances behavioral and molecular vulnerability. Male C57BL/6J mice were assigned to Control, CRS, BC, or BC+CRS groups and exposed for 21 consecutive days, followed by behavioral testing and molecular analyses of plasma-depleted whole blood and stress-related brain regions. BC exposure alone induced depressive-like behavior, and the combined BC+CRS condition showed the most pronounced phenotype. These findings indicate that inhaled BC is sufficient to influence stress-relevant behavior and may heighten vulnerability under chronic stress. At the molecular level, BC shifted peripheral responses toward a stress- and inflammation-associated state with reduced plasticity-related signaling, whereas CRS preferentially engaged glucocorticoid-responsive regulation. Combined BC+CRS exposure further altered plasticity- and transcription-related regulatory programs in blood and stress-related brain regions, with prominent changes in the nucleus accumbens. These condition-dependent molecular patterns suggest that BC engages blood–brain stress-related pathways in a context- and region-specific manner. Together, these findings identify inhaled BC as a neurobehaviorally relevant environmental hazard.

Graphical summary.
Effects of black carbon exposure and chronic restraint stress on depressive-like behavior and blood–brain molecular responses. Inhaled black carbon (BC) exposure induced depressive-like behavioral alterations and reshaped molecular profiles in plasma-depleted whole blood and stress-related brain regions. Molecular changes were classified as BC-associated, CRS-associated, shared BC/CRS-associated, or most evident in the BC+CRS condition. mPFC, medial prefrontal cortex; NAc, nucleus accumbens.

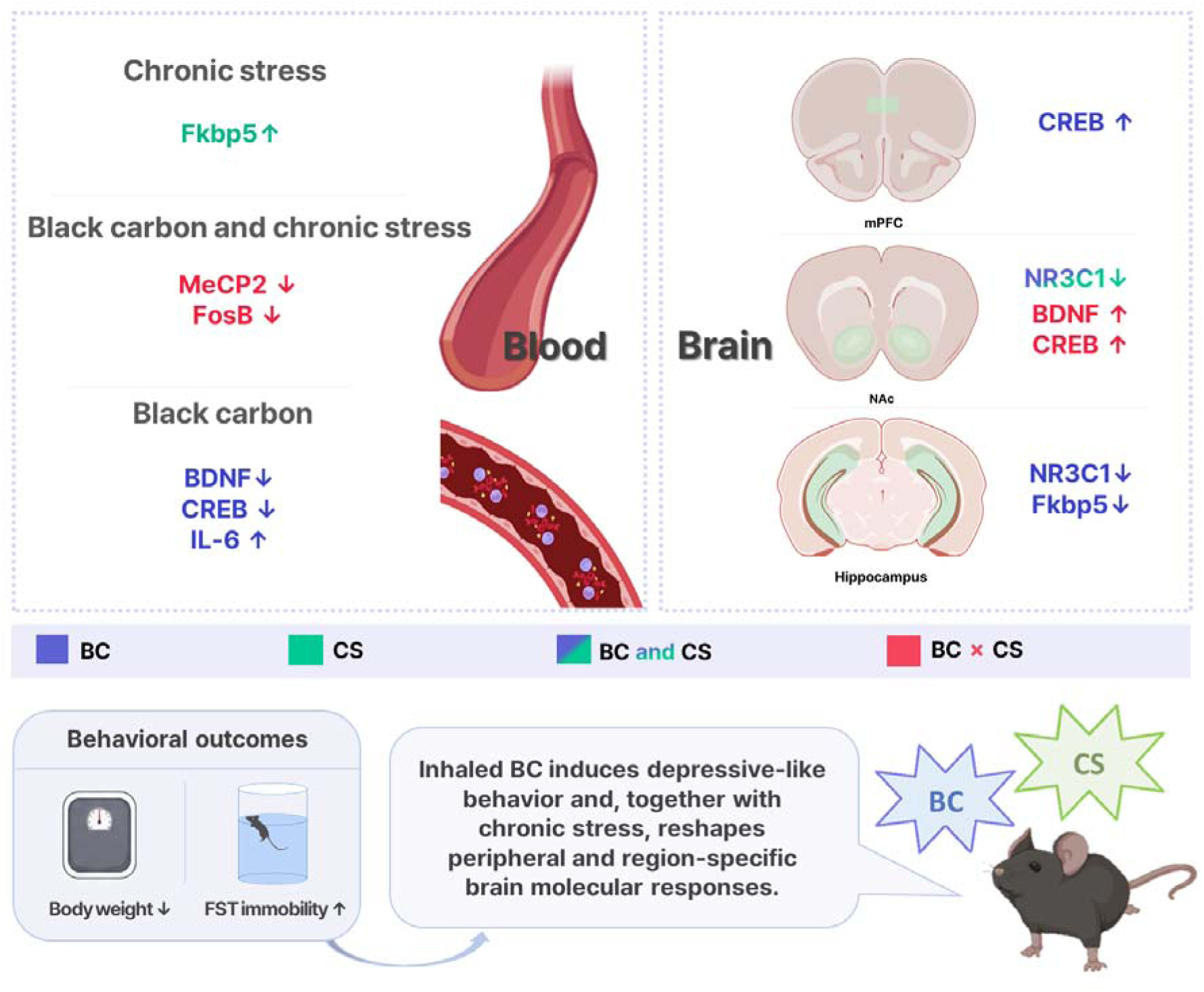

## Introduction

Black carbon (BC) is a major carbonaceous component of fine particulate matter (PM□.□) generated by incomplete combustion from common combustion-related sources, including diesel vehicle exhaust, residential heating, biomass burning, and industrial activity [1,2]. Because BC more specifically reflects combustion-derived particles than total PM mass, it has been proposed as an additional indicator of the adverse health effects of airborne particles compared with PM□□ and PM□.□ mass concentrations [3]. BC is widely recognized as an indicator of combustion-related air pollution and has been associated with cardiopulmonary morbidity and premature mortality [1-3]. Although BC-containing particulate matter has been studied mainly in relation to respiratory and cardiovascular outcomes, increasing attention has been directed toward its potential effects on brain function and mental health-related outcomes.

Recent epidemiological studies have reported associations between ambient BC or carbonaceous PM□.□ components and depressive symptoms or depression-related outcomes, with direct BC-related evidence emerging from studies of depressive symptoms and broader mental-health outcomes [4-6], and additional support from PM□.□ component-based cohort studies [7-10]. Systematic reviews and meta-analyses have further supported associations between particulate air pollution exposure and depression-related outcomes, while emphasizing heterogeneity among studies and the need for evidence addressing potential causal pathways [11,12]. A retrospective cohort study in college students reported that BC exposure was associated with symptoms of depression [5]. In addition, nationwide cohort studies in China found associations between PM□.□ components, including black carbon, and worsening depressive symptoms or depression among middle-aged and older adults [9,10]. A recent US Medicare cohort further reported that specific PM□.□ components, including elemental carbon, were associated with incident depression in older adults [7]. These findings raise the possibility that combustion-related PM components contribute to mood dysregulation. However, epidemiological studies remain largely correlational, and controlled experimental evidence is needed to determine whether inhaled BC exposure itself can induce depression-related behavioral alterations and to identify biological pathways linking BC exposure to stress-related mood vulnerability.

Experimental studies using PM exposure models suggest that inhaled particulate exposure can induce depressive-like behavioral alterations in rodents and affect inflammatory and BDNF-related signaling pathways [13,14]. In parallel, chronic stress engages endocrine, inflammatory, and neuroplasticity-related pathways, including glucocorticoid feedback signaling and activity-dependent plasticity-associated regulation [15-19]. Because real-world environmental exposure often occurs together with psychological stress, BC exposure may contribute not only to the onset of depressive-like behavioral alterations but also to increased vulnerability under chronic stress conditions. However, whether BC exposure alone is sufficient to induce depressive-like behavioral changes, and whether concurrent psychological stress reveals additional molecular alterations in peripheral and brain compartments, remains unclear.

In the present study, we established a controlled BC inhalation exposure model combined with chronic restraint stress (CRS) to examine the behavioral and molecular consequences of BC exposure in mice. We tested whether chronic BC exposure induces depressive-like behavioral alterations and whether combined BC+CRS exposure is associated with increased stress-related behavioral and molecular vulnerability. To define peripheral and central molecular responses, we examined stress-, inflammation-, and plasticity-related transcripts in plasma-depleted whole blood and representative stress-related brain regions, including the medial prefrontal cortex, nucleus accumbens, and hippocampus. This study provides preclinical evidence that inhaled BC exposure contributes to depression-related behavioral alterations and reshapes peripheral and brain-region-specific molecular responses under chronic stress conditions.

## Materials and methods

### Animals

Male C57BL/6J mice aged 7–9 weeks were obtained from Dae Han Bio Link Co., Ltd. (Eumseong, Republic of Korea). Mice were maintained under standard laboratory conditions at 21–23°C and 50–55% relative humidity under a 12-h light/dark cycle, with food and water provided ad libitum. Animals were randomly assigned to experimental groups in a 2 × 2 factorial design consisting of no black carbon exposure/no chronic restraint stress, black carbon exposure alone, chronic restraint stress alone, and combined black carbon exposure plus chronic restraint stress. All behavioral procedures were conducted during the light phase of the cycle. All animal experiments were approved by the Institutional Animal Care and Use Committee of the Brain Science Institute, Korea Institute of Science and Technology, and were performed in accordance with institutional guidelines.

### Experimental design

The study was designed to determine the independent and combined effects of inhaled black carbon exposure and chronic restraint stress on depression-related behavioral and molecular outcomes. Mice were assigned to four groups: Control, chronic restraint stress (CRS), black carbon exposure (BC), and combined BC plus CRS. BC exposure and CRS were applied over a 21-day experimental period. Body weight was monitored throughout the exposure period.

Behavioral tests were performed according to the experimental timeline, including the open field test, elevated plus maze test, and forced swim test. Blood, stress-related brain regions, and peripheral organs were collected approximately 4 days after the final exposure/restraint session for molecular analyses. Brain regions included the medial prefrontal cortex, nucleus accumbens, and hippocampus. Peripheral organs included the liver, lung, and distal colon.

### Black carbon generation and inhalation exposure

The BC generation method was adapted from a previously established soot generation system [20]. In this system, hydrocarbon fuels undergo pyrolysis, followed by nucleation and surface growth of primary soot particles and their subsequent agglomeration [20,21]. Such carbonaceous particles typically form aggregates of primary soot particles [21,22]. Previous work demonstrated that soot formation varies with pyrolyzer temperature, fuel mole fraction, and residence time in the furnace. Residence time, in particular, influences the extent of pyrolysis. Accordingly, the operating conditions used in this study were selected to promote pyrolysis and carbonaceous particle formation under controlled conditions [20].

BC aerosol was generated using a controlled pyrolysis particle generation system coupled to a whole-body inhalation exposure chamber. Propylene gas (C_3_H_6_) was used as the carbon source, and nitrogen gas (N_2_) was used as the carrier gas to maintain oxygen-deficient conditions in the pyrolyzer. Propylene and nitrogen were supplied at flow rates of 4 mL/min and 1 L/min, respectively, using independent mass flow controllers and were mixed in a gas mixer before entering the pyrolyzer. The gas mixture was then introduced into the pyrolyzer maintained at 1250 °C. Under these oxygen-deficient, high-temperature conditions, propylene underwent pyrolysis, leading to the formation and subsequent aggregation of primary carbonaceous particles.

All mice underwent a 6 h daily chamber session for 21 consecutive days. Mice in the control and CRS groups were placed in the control chamber, whereas mice in the BC and BC+CRS groups were placed in the BC exposure chamber. To prevent visual contact between CRS and non-CRS mice, a partition was installed between the corresponding sections within each chamber (Fig. 1a). Mice in the BC and BC+CRS groups underwent a 3 h BC exposure period within each 6 h chamber session. Mice in the control and BC groups remained unrestrained throughout the 6 h session, whereas mice in the CRS and BC+CRS groups underwent restraint for the entire 6 h session. Thus, in the BC+CRS group, the 3 h BC exposure occurred during the 6 h restraint period.

**Figure 1.**
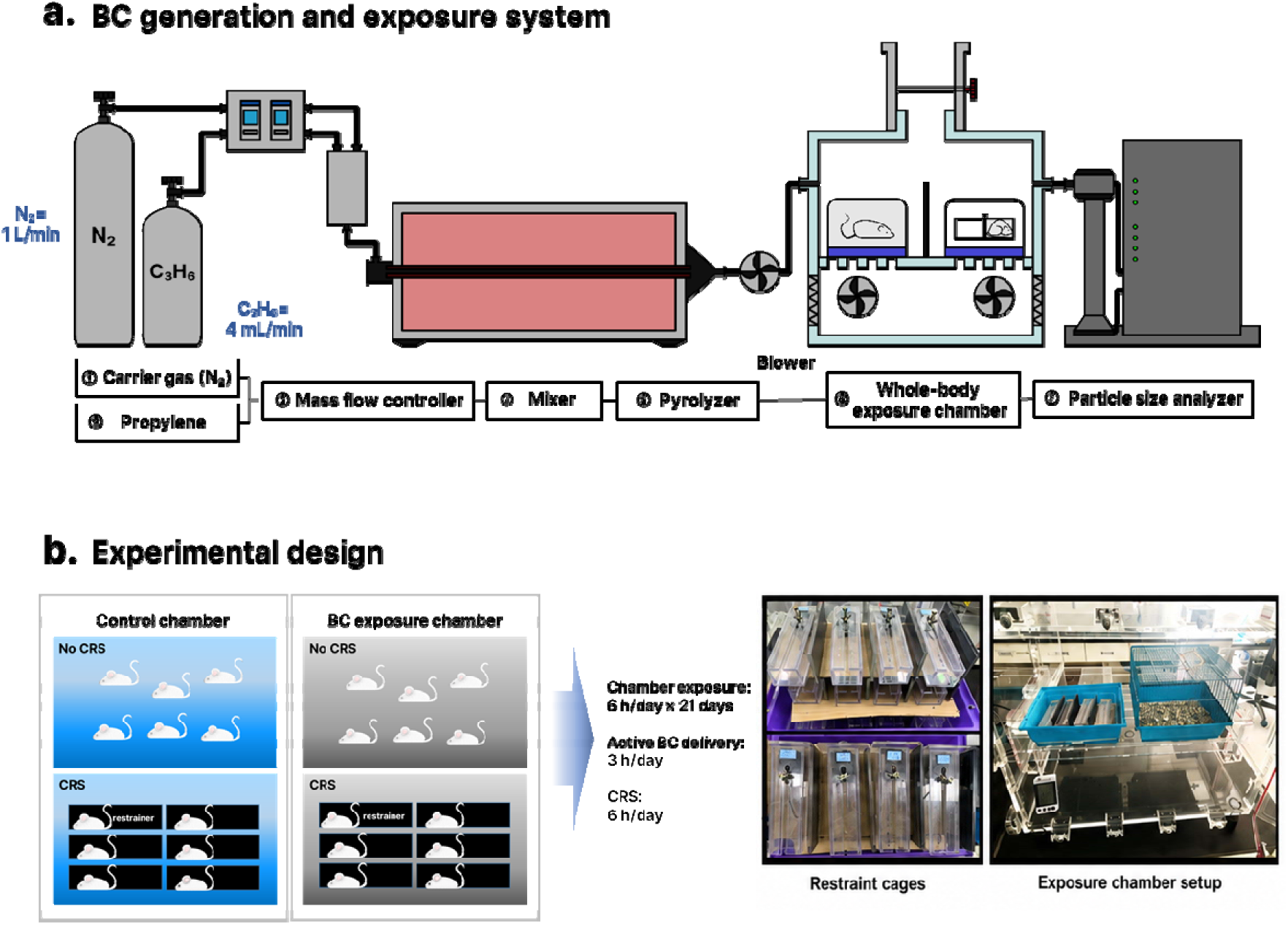
Overview of the black carbon generation and inhalation exposure system and experimental design. (a) Schematic illustration of the controlled black carbon (BC) generation and whole-body inhalation exposure system. Propylene gas (C□H□) was used as the carbon source, and nitrogen (N□) was used as the carrier gas. Gas flow was regulated using mass flow controllers, mixed, pyrolyzed at 1250°C, and delivered into the whole-body exposure chamber. Particle size distribution was monitored using a particle size analyzer. (b) Experimental design for chamber-matched BC exposure and chronic restraint stress (CRS). Mice were assigned to Control, CRS, BC, or BC+CRS groups. All mice were placed in the assigned chamber for 6 h/day for 21 consecutive days. Active BC aerosol delivery was performed for 3 h/day in the BC and BC+CRS groups, whereas CRS was applied for 6 h/day in the CRS and BC+CRS groups.

### Characterization of generated black carbon aerosol

BC exposure conditions were characterized by monitoring aerosol mass concentration, particle size distribution, and particle morphology in the exposure chamber. BC aerosol delivery was performed using a cyclic exposure protocol established in previous studies [23,24]. During the 3 h BC exposure period, seven exposure cycles were conducted, with ventilation between cycles to remove residual particles from the chamber. The exposure cycles were configured to maintain the chamber CO_2_ concentration below 2000 ppm. **Chamber aerosol mass concentration was monitored using a scanning mobility particle sizer equipped with a condensation particle counter (SMPS+C 5416, GRIMM Aerosol Technik, Germany) to characterize the temporal exposure profile. Aerosol mass concentration was automatically calculated by the instrument software. Measurements were acquired at 4 min intervals, corresponding to the measurement cycle of the instrument.**

**The particle number size distribution of the generated BC aerosol was determined using the SMPS+C 5416 to characterize particles in the submicrometer range.** Particle morphology and agglomeration were further examined by scanning electron microscopy (SEM; Regulus 8230, Hitachi High-Tech, Japan) at a magnification of 200,000×.

### Chronic restraint stress

Chronic restraint stress was performed using a rectangular restraint apparatus with an internal space of approximately 3 × 3 × 17 cm. Each mouse was placed individually into the restrainer, and the adjustable barrier was positioned to restrict forward and backward movement while allowing limited postural adjustment. Importantly, restraint stress was conducted inside the assigned exposure chamber. CRS-only mice were restrained in the control chamber, whereas BC+CRS mice were restrained in the black carbon exposure chamber. Mice in the CRS groups were restrained for 6 h per day for 21 consecutive days. Non-CRS mice were placed in their corresponding chamber without restraint during the exposure period. Routine cage maintenance and body weight monitoring were performed for all groups throughout the experiment.

### Behavioral tests

#### Body weight measurement

Body weight was measured repeatedly during the 21-day exposure and stress period. For longitudinal analysis, body weight values were normalized to each animal’s baseline body weight at the beginning of the experiment and expressed as a percentage of baseline. These data were used to evaluate the effects of BC exposure, CRS, and their combination on body weight trajectory across the experimental period.

### Open field test

Locomotor activity and center-zone exploration were assessed using the open field test. Mice were placed individually in a square open-field arena made of white acrylic material (40 × 40 × 40 cm) and allowed to freely explore the arena for 30 min. Mouse behavior was recorded using an overhead camera, and movement was analyzed using EthoVision XT software (Noldus Information Technology, Wageningen, The Netherlands). The center zone was defined as the central area of the arena (20 × 20 cm). Total distance moved and time spent in the center zone were used as behavioral readouts.

### Elevated plus maze

An elevated plus maze was used to assess anxiety-related behavior. The apparatus consisted of two open arms and two closed arms extending from a central platform. Closed arms were enclosed by walls, whereas open arms had no walls. The maze was elevated approximately 44 cm above the floor. Each mouse was placed in the central zone facing an open arm and allowed to explore the maze for 10 min. Behavior was recorded using a video camera. Time spent in the open arms was quantified, and arm entry was counted only when all four paws entered the arm. Time spent in the central platform was excluded from open- and closed-arm duration measurements.

### Forced swim test

Depressive-like behavior was evaluated using the forced swim test. Mice were individually placed in a transparent cylinder filled with water maintained at 24 ± 1°C. The test lasted 6 min, and immobility time was analyzed during the final 4 min. Fresh water was used for each animal. Immobility was defined as the absence of active escape-directed movement, with only minimal movements required to keep the head above water. Immobility duration was scored manually by an experimenter blinded to group.

### Blood collection and plasma-depleted whole blood preparation

Whole blood was collected into heparin-coated tubes and kept on ice until processing. Samples were centrifuged at 300 × g for 10 min at 4°C to separate the plasma from the cellular fraction. After centrifugation, the plasma supernatant was carefully removed by pipetting without disturbing the lower cellular fraction. The remaining plasma-depleted whole blood fraction, containing erythrocytes and residual leukocytes, was immediately lysed in TRIzol™ LS reagent. Total RNA was extracted according to the manufacturer’s instructions. RNA quantity and purity were assessed by spectrophotometry, and samples with acceptable A260/A280 ratios were used for cDNA synthesis and quantitative PCR analysis. Detection of *Cd3e* transcripts confirmed the presence of residual leukocytes in the plasma-depleted whole blood fraction used for RNA analysis.

### Plasma corticosterone measurement

Plasma corticosterone concentrations were determined using a Corticosterone Competitive ELISA Kit (Invitrogen, EIACORT). Plasma samples were diluted 1:200 with Dissociation Reagent and 1× Assay Buffer. Corticosterone standards ranging from 0 to 10,000 pg/mL and diluted plasma samples were loaded in duplicate at 50 µL per well. Corticosterone Conjugate and Corticosterone Antibody were added to each well at 25 µL per well, except for the non-specific binding wells. Plates were incubated for 1 h at room temperature with shaking and washed four times with 1× Wash Buffer. TMB Substrate was then added and incubated for 30 min, followed by Stop Solution. Absorbance was measured at 450 nm using a microplate reader. Corticosterone concentrations were calculated from a four-parameter logistic standard curve and corrected for the dilution factor. Corticosterone values were normalized to the mean value of the Control group and are presented as relative plasma corticosterone levels.

### Tissue collection and brain-region dissection

Mice were sacrificed approximately 4 days after the final BC exposure and/or restraint stress session. Brains were rapidly removed and placed on an ice-cold surface. The medial prefrontal cortex, nucleus accumbens, and hippocampus were dissected according to anatomical landmarks based on the mouse brain atlas. Dissected tissues were immediately frozen on dry ice or in liquid nitrogen and stored at −80°C until RNA extraction. All brain dissections were performed under consistent conditions to minimize tissue degradation and regional variability.

### Peripheral organ collection

Peripheral organs were collected to examine organ-specific inflammatory transcript responses. Liver, lung, and distal colon tissues were rapidly dissected after sacrifice. For colon samples, the distal segment was isolated, and luminal contents were gently removed by rinsing with cold PBS. Tissue samples were immediately frozen on dry ice or in liquid nitrogen and stored at −80°C until RNA extraction. These samples were used for exploratory analysis of inflammatory transcripts, including *Il6*, *Il1b*, and *Tnf*.

### RNA extraction and cDNA synthesis

Total RNA was extracted from dissected brain regions and peripheral organ tissues using TRIzol reagent, and from plasma-depleted whole blood using TRIzol LS reagent, according to the manufacturer’s instructions. Frozen tissue samples were thoroughly homogenized in TRIzol reagent before phase separation and RNA precipitation. RNA concentration and purity were assessed using NanoDrop. Complementary DNA was synthesized from total RNA using ReverTra Ace qPCR RT Master Mix (TOYOBO, Osaka, Japan) following the manufacturer’s protocol. The synthesized cDNA was diluted as appropriate and used for quantitative real-time PCR.

### Quantitative real-time PCR

Quantitative real-time PCR was performed to measure inflammatory, glucocorticoid-related, neuroplasticity-related, and transcription-related genes in plasma-depleted whole blood, dissected brain regions, and peripheral organ tissues. Quantitative PCR was performed using THUNDERBIRD SYBR qPCR Master Mix (TOYOBO, Osaka, Japan) on a CFX Connect Real-Time PCR Detection System (Bio-Rad, CA, USA).

The same primer pairs were used for each target transcript across different tissue types. Relative mRNA expression levels were calculated using the ΔΔCt method, with Gapdh used as the internal reference gene. Expression values were normalized to the mean value of the control group and are presented as relative mRNA expression fold change. Primer sequences used in this study are listed in Supplementary Table S1.

### Detection of residual leukocyte-associated transcripts

To confirm that plasma-depleted whole blood samples contained residual leukocyte-associated RNA, *Cd3e* expression was examined by quantitative real-time PCR. This analysis was used as a qualitative confirmation of leukocyte-associated transcript detection, and blood molecular data were interpreted as transcript profiles from plasma-depleted whole blood rather than purified PBMCs. Primer sequences are listed in Supplementary Table S1.

### Statistical analysis

Data are presented as mean ± SEM. Statistical analyses were performed using GraphPad Prism software. Endpoint behavioral and molecular data were analyzed using two-way ANOVA to assess the main effects of BC exposure and CRS, and their interaction. When appropriate, pairwise group comparisons were performed using Fisher’s least significant difference post hoc test. Body weight data collected over time were analyzed using repeated-measures two-way ANOVA, followed by Fisher’s least significant difference post hoc test when appropriate. Statistical significance was set at *p* < 0.05. In the figures, asterisks denote significant differences from the Control group, number signs denote significant differences from the CRS group, and dollar signs denote significant differences from the BC group, where indicated.

## Results

### Experimental design of controlled BC inhalation exposure combined with chronic restraint stress

A controlled BC inhalation exposure paradigm combined with chronic restraint stress (CRS) was used to examine the independent and combined effects of BC and CRS (Fig. 1). Mice were assigned to four experimental groups: Control, CRS, BC, and BC+CRS. All mice underwent a 6 h daily chamber session for 21 consecutive days. Control and CRS mice were placed in the control chamber, whereas BC and BC+CRS mice were placed in the BC exposure chamber. To prevent visual contact between CRS and non-CRS mice, a partition was installed between the corresponding sections of each chamber (Fig. 1a).

Mice in the BC and BC+CRS groups underwent a 3 h BC exposure period within each 6 h chamber session. CRS was applied throughout the entire 6 h session in the CRS and BC+CRS groups. Thus, mice in the BC group remained unrestrained throughout the 6 h chamber session, whereas mice in the BC+CRS group underwent 6 h of restraint, including the 3 h BC exposure period (Fig. 1b). **Behavioral testing was performed after the 21-day exposure period, and blood and tissue samples were collected on post-exposure day 4 for molecular analyses.**

### Characterization of BC generation and inhalation exposure conditions

We next characterized the BC exposure environment generated by the pyrolysis-based inhalation system. During each 3 h BC exposure period, chamber aerosol mass concentration showed a reproducible cyclic pattern across seven exposure cycles (Fig. 2a). Aerosol mass concentration increased at the beginning of each cycle and gradually declined thereafter. This cyclic exposure pattern was consistent with the chamber exposure approach established in previous studies [23,24].

**Figure 2.**
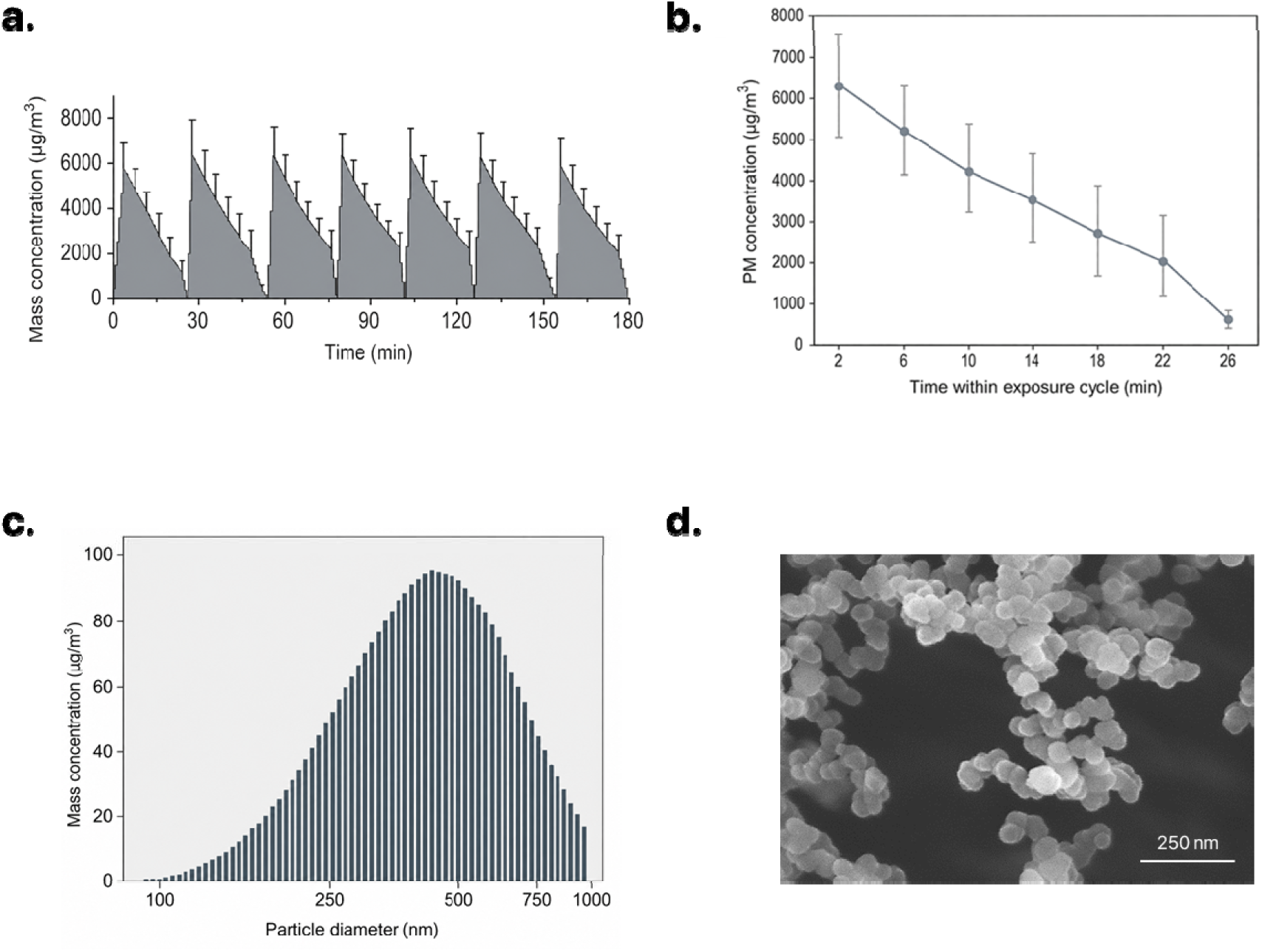
Characterization of black carbon exposure. (a) Repeated chamber aerosol mass concentration profiles during the 3-h active black carbon (BC) delivery period. (b) Average chamber aerosol mass concentration profile within each 30-min BC exposure cycle. Data in (a) and (b) are presented as mean ± SEM. (c) The mass-based particle size distribution of the generated BC aerosol was determined using the SMPS+C 5416. The generated particles were predominantly distributed in the submicrometer range, with most falling within the PM□ size range. (d) Representative scanning electron microscopy image of generated BC particles showing aggregated carbonaceous particle morphology. Scale bar, 250 nm.

When the concentration profiles were averaged across exposure cycles, the chamber aerosol mass concentration was initially approximately 6 × 10³ µg/m³ and progressively declined over the course of each cycle (Fig. 2b). Particle size distribution analysis showed that the generated aerosol was predominantly composed of submicrometer particles (Fig. 2c). Scanning electron microscopy further revealed aggregated carbonaceous particles (Fig. 2d), consistent with the morphology previously reported for pyrolysis-generated soot [20]. Together, these results demonstrate that the pyrolysis-based exposure system generated and delivered BC aerosol with a consistent cyclic exposure profile under controlled chamber conditions.

Using these characterized exposure conditions, we next examined the behavioral effects of BC exposure alone and in combination with CRS.

### BC exposure and CRS induce depressive-like behavioral alterations

After 21 days of BC exposure and/or CRS, mice underwent behavioral testing followed by blood and tissue collection (Fig. 3a). Body weight gain was reduced by BC exposure and CRS, with the strongest reduction observed in the BC+CRS group (Fig. 3b). To determine whether these treatments affected general locomotor activity, mice were first examined in the open field test. Total distance moved and center-zone duration were not significantly changed across groups, indicating that BC exposure and CRS did not produce major locomotor impairment (Fig. 3c–e).

**Figure 3.**
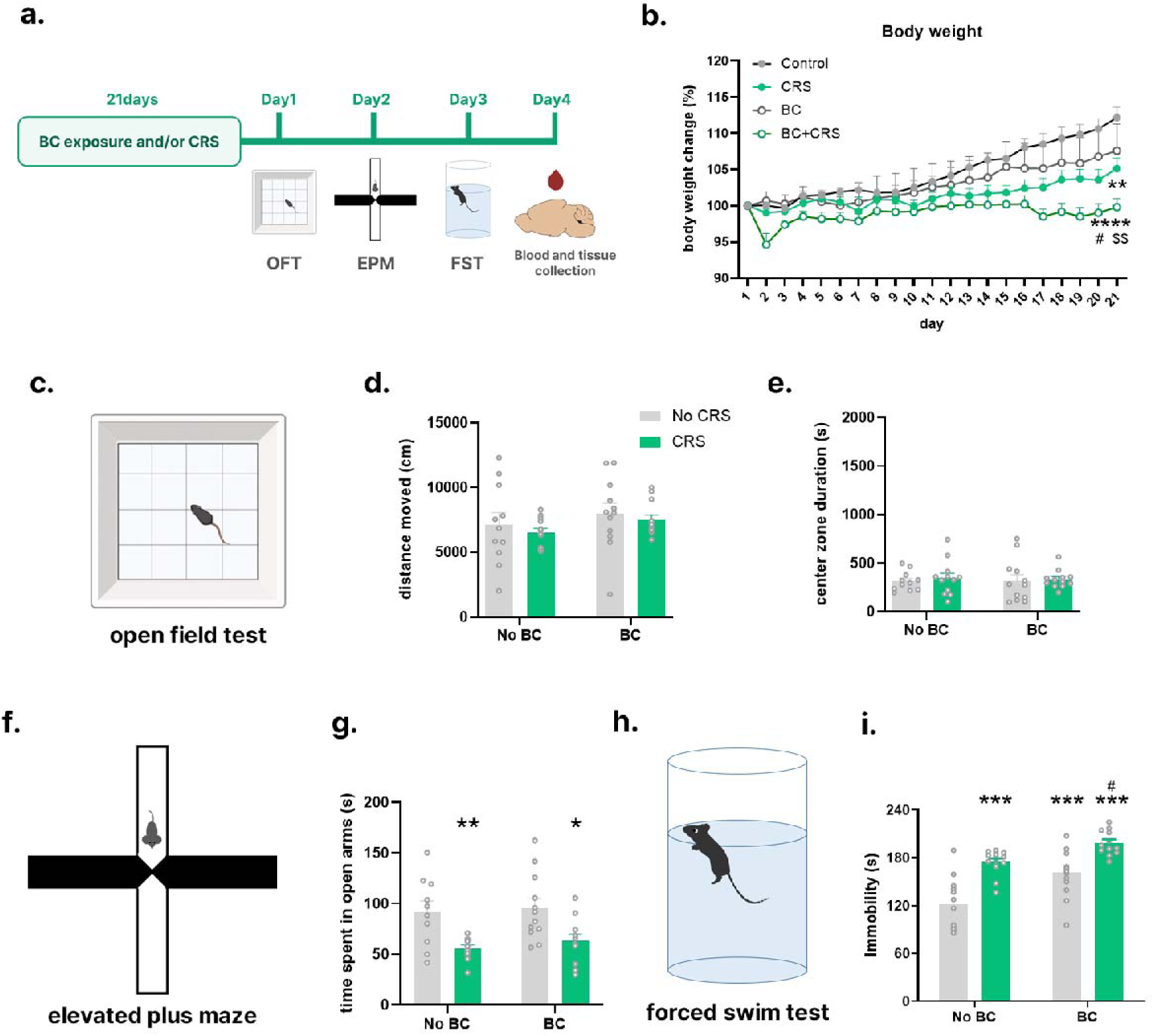
Behavioral alterations following black carbon exposure and chronic restraint stress. (a) Experimental timeline for BC exposure, CRS, behavioral testing, and tissue collection. Mice were exposed to BC and/or CRS for 21 days, followed by open field test (OFT), elevated plus maze (EPM), forced swim test (FST), and blood and tissue collection over four consecutive days. (b) Body weight changes during the experimental period. (c) Schematic illustration of the OFT. (d, e) Total distance traveled and center-zone activity in the OFT. (f) Schematic illustration of the EPM. (g) Open-arm activity in the EPM. (h) Schematic illustration of the FST. (i) Immobility time in the FST. BC exposure and CRS increased immobility in the FST, without a significant BC × CRS interaction. OFT activity was not significantly affected by BC exposure, CRS, or their interaction. EPM behavior was affected by CRS, but not by BC exposure or the BC × CRS interaction. Data are presented as mean ± SEM. Statistical details are described in Methods. n = 10–12 mice per group. n.s., not significant; *p < 0.05, **p < 0.01, ***p < 0.001 vs. Control; #p < 0.05 vs. CRS.

Anxiety-like behavior was then assessed using the elevated plus maze. CRS reduced the time spent in the open arms, whereas BC exposure alone did not show a clear effect on this measure (Fig. 3f, g). These results suggest that anxiety-like behavioral alteration was mainly associated with CRS under the present experimental condition. In contrast, depressive-like behavior assessed by the forced swim test was affected by both BC exposure and CRS. Immobility time was increased in both BC-exposed and CRS-exposed mice, and the BC+CRS group showed the highest immobility level among the four groups (Fig. 3h, i). These findings indicate that controlled BC exposure is sufficient to induce depressive-like behavior, while the combined BC+CRS condition produces the most pronounced depressive-like behavioral outcome.

### BC exposure and CRS produce distinct peripheral molecular responses in plasma-depleted whole blood

Because BC exposure and CRS produced depressive-like behavioral alterations, we next examined whether these conditions were accompanied by systemic molecular changes in plasma-depleted whole blood (Fig. 4). We first assessed corticosterone and glucocorticoid-related transcripts to evaluate peripheral stress-related responses. Relative plasma corticosterone levels were increased following BC exposure and CRS, suggesting that both inhaled BC exposure and chronic restraint stress engaged systemic endocrine stress-related responses (Fig. 4a). Among glucocorticoid-related transcripts, *Fkbp5* expression was increased in CRS-exposed mice, consistent with prior evidence that *Fkbp5* is a glucocorticoid-responsive stress-related transcript [25] (Fig. 4b). In contrast, blood *Nr3c1* expression was not significantly altered (Supplementary Fig. S1), indicating that the peripheral stress response was reflected more prominently at the level of circulating corticosterone and downstream glucocorticoid-responsive transcription than at the level of glucocorticoid receptor transcript abundance.

**Figure 4.**
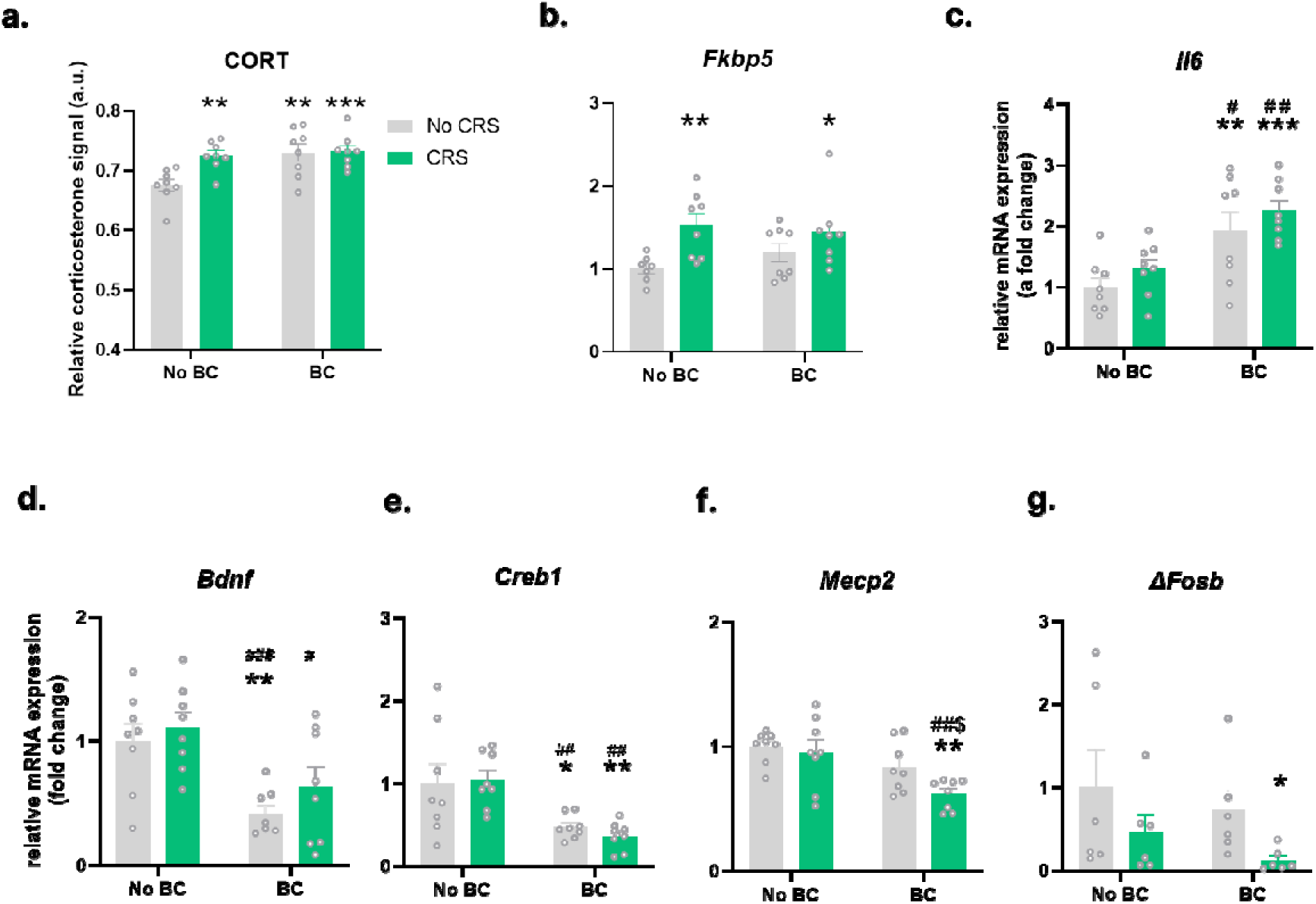
Distinct peripheral molecular responses to black carbon exposure and chronic restraint stress. (a) Relative corticosterone signal (a.u.) (b–g) Relative mRNA expression of *Fkbp5*, *Il6*, *Bdnf*, *Creb1*, *Mecp2*, and ΔFosB-related *Fosb* transcript in plasma-depleted whole blood. Corticosterone was increased following BC exposure and CRS. *Fkbp5* showed a CRS-associated increase, whereas *Il6*, *Bdnf*, and *Creb1* showed BC-associated changes. *Mecp2* and ΔFosB-related *Fosb* transcript were reduced most clearly in the BC+CRS group. Data are presented as mean ± SEM. Statistical details are described in Methods. n = 6–8 mice per group. n.s., not significant; *p < 0.05, **p < 0.01, ***p < 0.001 vs. Control; #p < 0.05, ##p < 0.01, ###p < 0.001 vs. CRS; $p < 0.05 vs. BC.

Because inhaled particulate exposure can affect immune and inflammatory signaling, we next examined inflammatory transcripts in blood. *Il6* expression was elevated in BC-exposed groups, indicating that BC exposure engaged an inflammatory component of the peripheral response (Fig. 4c). Additional inflammatory transcripts, including *Il1b* and *Tnf*, were examined as supplementary markers and did not show significant changes in plasma-depleted whole blood (Supplementary Fig. S1). We then extended this analysis to peripheral organs to determine whether inflammatory transcript changes were also detectable outside the blood compartment. In the liver, inflammatory transcript expression was generally reduced, whereas lung inflammatory transcripts showed limited changes. In the colon, *Il6* was elevated most prominently in the BC+CRS group, and *Il1b* was increased in BC-exposed groups (Supplementary Fig. S2). These data indicate that BC exposure and CRS were accompanied by compartment-dependent peripheral inflammatory responses involving blood and selected peripheral tissues.

We next asked whether these endocrine and inflammatory peripheral responses were accompanied by changes in molecular regulators linked to stress adaptation, neuroplasticity, synaptic regulation, and depression-related behavioral outcomes. *Bdnf* and *Creb1* were examined because BDNF–CREB signaling is a central activity-dependent pathway involved in neuronal plasticity, stress adaptation, and affective regulation [15]. In plasma-depleted whole blood, BC exposure was associated with reduced *Bdnf* and *Creb1* expression (Fig. 4d, e), suggesting that BC exposure was accompanied by peripheral changes in plasticity-related regulatory markers.

We also examined *Mecp2* and ΔFosB-related *Fosb* transcript as depression-relevant transcriptional regulators linked to activity-dependent and synaptic plasticity-related molecular adaptation [26,27]. In the present study, *Mecp2* expression was reduced most clearly in the BC+CRS group, and ΔFosB-related *Fosb* transcript showed a similar reduction under the combined BC+CRS condition (Fig. 4f, g). Notably, these reductions were not evident in either single-exposure group but emerged most clearly when BC exposure and CRS were combined, suggesting that concurrent environmental and psychological stressors were associated with a more pronounced reduced expression of depression-relevant plasticity-related transcriptional regulators.

Together, these results indicate that BC exposure and CRS were accompanied by layered peripheral molecular alterations. CRS was linked to endocrine and glucocorticoid-responsive stress signaling, BC exposure was associated with inflammatory and plasticity-related transcript changes, and the combined BC+CRS condition revealed reduced expression of depression-relevant transcriptional regulators, including *Mecp2* and ΔFosB-related *Fosb* transcript.

### BC exposure and CRS induce region-specific molecular responses in representative stress-related brain regions

Peripheral molecular changes provide systemic readouts of BC exposure and CRS, but depressive-like behavioral outcomes are likely mediated through distributed stress-related brain circuits. Therefore, we next examined molecular responses in three representative stress-related brain regions—the mPFC, NAc, and hippocampus—which are closely involved in affective regulation, stress adaptation, and depression-related neuroplasticity [28,29].

In the mPFC, *Creb1* expression was increased following BC exposure (Fig. 5a, b). ΔFosB-related *Fosb* transcript showed a condition-dependent expression pattern, with increased expression in the CRS-only group and a relative reduction when BC exposure was combined with CRS (Fig. 5c). Given the roles of CREB and ΔFosB in activity-dependent transcriptional regulation and plasticity-related molecular adaptation, these findings suggest that BC exposure and CRS affected transcriptional regulatory markers in the mPFC in a condition-dependent manner [15,27].

**Figure 5.**
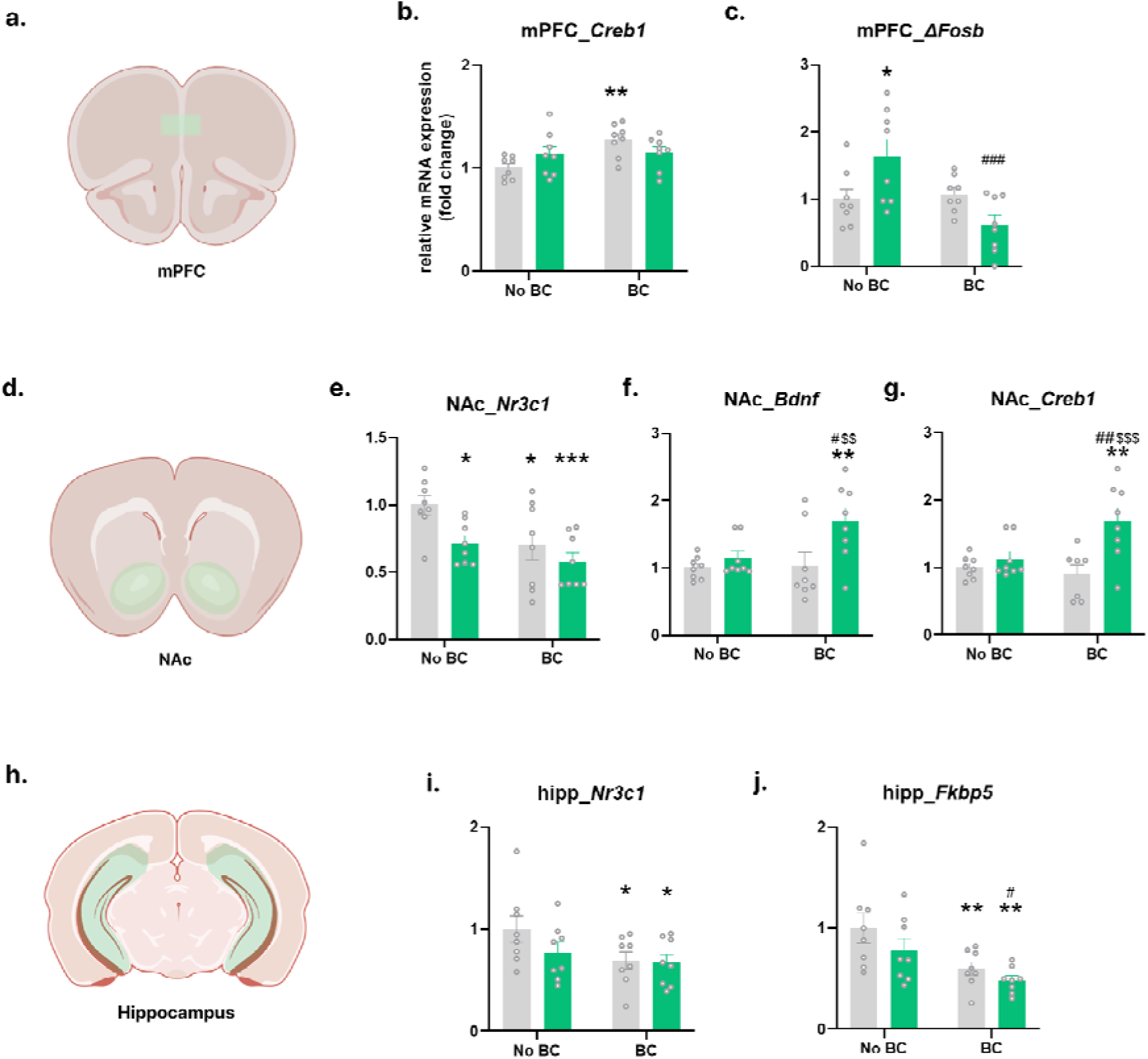
Region-specific brain molecular responses to black carbon exposure and chronic restraint stress. (a) Schematic illustration of the medial prefrontal cortex (mPFC). (b, c) Relative mRNA expression of *Creb1* and ΔFosB-related *Fosb* transcript in the mPFC. (d) Schematic illustration of the nucleus accumbens (NAc). (e–g) Relative mRNA expression of *Nr3c1*, *Bdnf*, and *Creb1* in the NAc. (h) Schematic illustration of the hippocampus. (i, j) Relative mRNA expression of *Nr3c1* and *Fkbp5* in the hippocampus. In the mPFC, *Creb1* and ΔFosB-related *Fosb* transcript showed BC-associated changes with BC × CRS-dependent patterns. In the NAc, *Nr3c1* was reduced by BC exposure and CRS, whereas *Bdnf* and *Creb1* were elevated in the BC+CRS group. In the hippocampus, *Fkbp5* was reduced by BC exposure, while *Nr3c1* showed modest reductions in BC-exposed groups. Data are presented as mean ± SEM. n = 7–8 mice per group. n.s., not significant; *p < 0.05, **p < 0.01, ***p < 0.001 vs. Control; #p < 0.05, ##p < 0.01, ###p < 0.001 vs. CRS; $$p < 0.01, $$$p < 0.001 vs. BC.

We then examined the NAc, a representative region involved in motivation, reward processing, stress susceptibility, and depressive-like behavioral regulation [28]. In the NAc, *Nr3c1* expression was reduced following both BC exposure and CRS (Fig. 5d, e), indicating that environmental and psychological stressors converged on glucocorticoid receptor-related transcriptional regulation in this region. In contrast, *Bdnf* and *Creb1* expression were increased most prominently in the BC+CRS group (Fig. 5f, g). These results suggest that, among the examined brain regions and markers, the combined BC+CRS condition produced the clearest plasticity-related transcriptional response in the NAc.

Finally, we examined the hippocampus, a representative stress-sensitive region involved in glucocorticoid feedback regulation, HPA-axis modulation, and stress-related affective adaptation [16,29]. In the hippocampus, *Nr3c1* and *Fkbp5* expression were reduced in BC-exposed groups, with the decrease in *Fkbp5* showing a clearer BC-associated pattern (Fig. 5h–j). Given the role of the hippocampus in glucocorticoid feedback control, these findings suggest that BC exposure affected hippocampal stress-regulatory transcriptional markers in a manner closely aligned with the functional role of this region. Thus, within the markers examined, the hippocampal response was most evident in glucocorticoid-related transcripts and appeared more closely associated with BC exposure than with CRS.

Additional screened markers across the mPFC, NAc, and hippocampus are summarized in Supplementary Fig. S3, and exploratory inflammatory transcript analyses are shown in Supplementary Fig. S4.

Together, these brain-region analyses indicate that BC exposure and CRS produced region-dependent molecular responses across representative stress-related brain regions. Notably, the NAc showed the clearest molecular signature of combined BC+CRS exposure, with *Bdnf* and *Creb1* most prominently increased in the BC+CRS group. In contrast, the hippocampus showed BC-associated reductions in glucocorticoid-related transcripts, consistent with its role in HPA-axis feedback regulation, whereas the mPFC showed changes in activity-dependent transcriptional regulatory markers. These findings suggest that concurrent BC exposure and CRS were most closely associated with plasticity-related molecular regulation in the NAc, while BC exposure was more closely linked to hippocampal glucocorticoid-regulatory changes.

### Integrated summary of BC- and CRS-associated blood–brain molecular alterations

To integrate the peripheral and brain molecular findings, we summarized the dominant molecular patterns observed in plasma-depleted whole blood and representative stress-related brain regions (Fig. 6). BC-associated changes included increased blood CORT and *Il6*, reduced blood *Bdnf* and *Creb1*, increased mPFC *Creb1*, and reduced hippocampal glucocorticoid-related transcripts. CRS-associated changes included increased blood *Fkbp5* and reduced NAc *Nr3c1*. Notably, the combined BC+CRS condition revealed the most prominent depression-relevant molecular alterations, including reduced blood *Mecp2* and ΔFosB-related *Fosb* transcript and increased NAc *Bdnf* and *Creb1*. These findings support a model in which inhaled BC exposure induces depressive-like behavior and systemic molecular changes, while concurrent CRS further reshapes selected plasticity- and transcription-related responses in a compartment- and brain-region-dependent manner.

**Figure 6.**
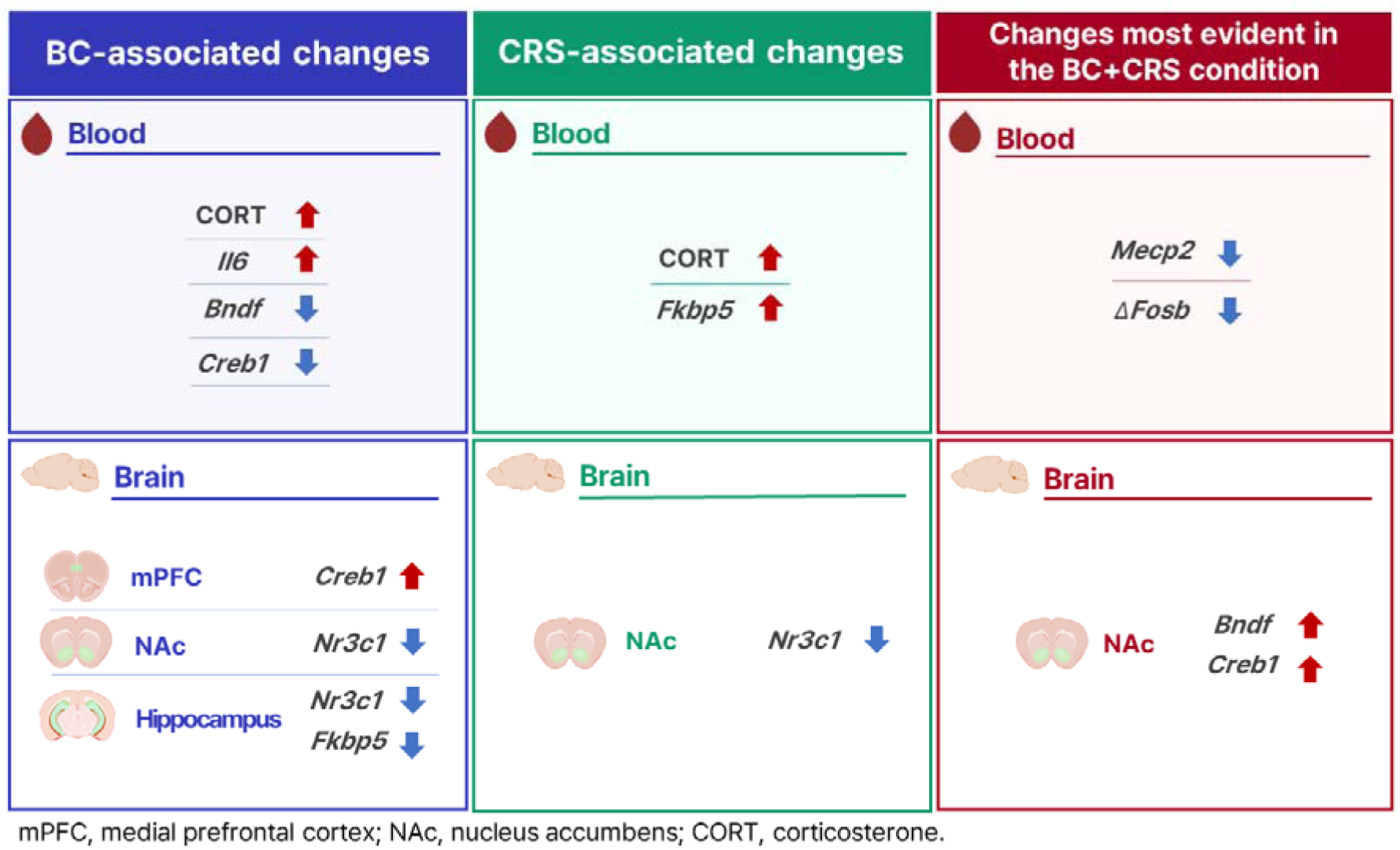
Summary of molecular alterations in blood and brain induced by black carbon exposure and chronic restraint stress. Schematic summary of molecular changes in plasma-depleted whole blood and stress-related brain regions following BC exposure, CRS, or combined BC+CRS exposure. Molecular alterations are grouped according to the dominant significant patterns observed in Figs. 4 and 5, including BC-associated changes, CRS-associated changes, and changes most evident in the BC+CRS condition. In blood, BC-associated changes included increased CORT and *Il6* and decreased *Bdnf* and *Creb1*, whereas CRS-associated changes included increased CORT and *Fkbp5*. *Mecp2* and ΔFosB-related *Fosb* transcript showed reductions most evident in the BC+CRS condition. In the brain, BC-associated changes included increased mPFC *Creb1*, decreased NAc *Nr3c1*, and decreased hippocampal *Nr3c1* and *Fkbp5*. CRS-associated brain changes included decreased NAc *Nr3c1*. Changes most evident in the BC+CRS condition were observed in the NAc, where *Bdnf* and *Creb1* were increased. mPFC, medial prefrontal cortex; NAc, nucleus accumbens; CORT, corticosterone. Italicized gene symbols indicate mRNA transcripts measured by qPCR. Arrows indicate the direction of significant changes shown in Figs. 4 and 5.

## Discussion

In this study, we established a controlled BC inhalation exposure model combined with CRS to determine whether inhaled BC contributes to depression-related behavioral and molecular alterations. A central finding of this study is that BC exposure alone induced depressive-like behavioral alterations, providing preclinical evidence that inhaled BC can contribute to depression-related outcomes. Importantly, BC exposure did not act only as an independent environmental stressor. The combined BC+CRS condition was associated with a more pronounced depressive-like behavioral phenotype and selected molecular alterations in depression-relevant plasticity- and transcription-related regulators, including reduced blood *Mecp2* and ΔFosB-related *Fosb* transcript and increased NAc *Bdnf* and *Creb1*. These findings suggest that BC exposure may both induce depressive-like behavioral alterations and increase stress-related behavioral and molecular vulnerability under chronic stress conditions.

Epidemiological and meta-analytic evidence has linked particulate air pollution exposure with depressive symptoms, depression, and related mental-health outcomes [11,12]. Recent component-based studies further suggest that BC or carbonaceous PM□.□ components are associated with depressive symptoms and depression-related outcomes [4-8,10]. However, population-based studies cannot fully determine whether BC-containing particulate exposure directly contributes to depression-related biological changes. Our controlled exposure model addresses this gap by showing that BC exposure alone induced depressive-like behavioral alterations in mice. In addition, the combined BC+CRS condition showed a more pronounced depressive-like behavioral phenotype, suggesting that BC exposure may also worsen stress-related depressive-like outcomes under chronic stress conditions. Thus, our findings provide experimental support for the biological plausibility of the association between BC exposure and depression-related outcomes.

Although the present study did not use radiolabeled particles or directly quantify tissue deposition of BC, recent quantitative particle-tracing work demonstrated that inhaled carbonaceous PM can undergo systemic translocation beyond the lung and be detected in peripheral organs and brain tissue [23]. This prior evidence supports the rationale for examining molecular alterations in plasma-depleted whole blood and representative stress-related brain regions after controlled BC inhalation exposure.

The peripheral molecular data indicate that BC exposure and CRS engaged partially distinct stress-related biological pathways. CORT was increased following both BC exposure and CRS, suggesting that inhaled environmental exposure can activate systemic endocrine stress-related responses. Among glucocorticoid-related transcripts, *Fkbp5* was increased mainly under CRS conditions, whereas blood *Nr3c1* expression was not significantly altered. Because *Fkbp5* is a glucocorticoid-responsive transcript involved in stress hormone feedback regulation [25], this pattern suggests that the peripheral glucocorticoid-related response was reflected more clearly by circulating CORT and downstream stress-responsive transcription than by changes in glucocorticoid receptor transcript abundance. In parallel with these endocrine stress-related changes, BC exposure increased blood *Il6*, whereas *Il1b* and *Tnf* were not significantly altered in plasma-depleted whole blood. Given that IL-6-related inflammatory signaling has been implicated in stress- and depression-related biological responses [17,18], the selective increase in blood *Il6* suggests that BC exposure engaged an inflammatory component of the peripheral stress response. Together with the peripheral organ data, including elevated colonic *Il6* and *Il1b* expression and limited inflammatory changes in the lung, these findings indicate that BC exposure induced compartment-specific inflammatory alterations rather than a uniform systemic cytokine response.

In addition to endocrine and inflammatory responses, BC exposure was associated with reduced blood *Bdnf* and *Creb1*. BDNF–CREB signaling is closely linked to activity-dependent plasticity, stress adaptation, and affective regulation [15,19,30]. Therefore, the reduction of these transcripts suggests that BC exposure may influence peripheral molecular readouts related to depression-relevant plasticity-associated regulation. Although blood transcript levels do not directly indicate synaptic changes in the brain, they may reflect broader stress-related biological adaptation accompanying depressive-like behavior. This interpretation is consistent with prior studies linking particulate exposure, inflammation, and altered BDNF/CREB-related signaling in stress- or depression-related models [13,14].

A notable finding was that *Mecp2* and ΔFosB-related *Fosb* transcript levels were reduced most clearly in the combined BC+CRS group. MeCP2 and ΔFosB are transcriptional regulators implicated in neuronal adaptation, synaptic plasticity, and stress-related behavioral phenotypes [26,27]. MeCP2 is particularly relevant because our recent work identified MeCP2 as a synaptic plasticity-related regulator of chronic stress-induced depressive-like behavior [26]. In the present study, the reductions in *Mecp2* and ΔFosB-related *Fosb* transcript were not evident in either single-exposure group but emerged most clearly when BC exposure and CRS were combined. This suggests that concurrent environmental and psychological stressors may be associated with a more pronounced reduction of depression-relevant plasticity-related transcriptional regulators in the periphery. We do not interpret this as evidence for a generalized synergistic interaction, but rather as an indication that combined exposure can reveal selected molecular alterations that are less apparent under either stressor alone.

The brain-region analysis further showed that BC exposure and CRS did not produce a uniform molecular response across representative stress-related brain regions. Among the examined markers and regions, the NAc showed the clearest molecular signature of combined BC+CRS exposure, with *Bdnf* and *Creb1* most prominently increased in the BC+CRS group. The NAc is a stress-sensitive reward-related region implicated in motivation, stress susceptibility, and depressive-like behavioral regulation [28,31,32]. Therefore, the increase in NAc *Bdnf* and *Creb1* may reflect a plasticity-related molecular response to the combined environmental and psychological stress condition. Interestingly, this direction differed from the blood, where BC exposure reduced *Bdnf* and *Creb1*, indicating that peripheral and brain molecular responses do not simply mirror each other. Instead, BC exposure and CRS appear to reshape depression-relevant molecular pathways in a compartment- and region-dependent manner.

The hippocampal response showed a different pattern. The hippocampus is a major stress-sensitive region involved in glucocorticoid feedback regulation and HPA-axis modulation [16,29,33,34]. In this region, *Nr3c1* and *Fkbp5* were reduced in BC-exposed groups, with *Fkbp5* showing a clearer BC-associated pattern. Because *Nr3c1* encodes the glucocorticoid receptor and *Fkbp5* is involved in glucocorticoid receptor sensitivity and feedback regulation [25,35,36], these changes align with the known role of the hippocampus in stress hormone regulation. Thus, whereas the NAc showed the most evident response to combined BC+CRS exposure in plasticity-related markers, the hippocampus showed BC-associated alterations in glucocorticoid-regulatory transcripts. This regional distinction supports the idea that BC exposure affects multiple stress-related brain systems through functionally different molecular routes.

Together, the behavioral and molecular findings suggest that BC exposure may contribute to depression-related vulnerability in two ways. First, BC exposure alone was sufficient to induce depressive-like behavioral alterations and peripheral molecular changes, supporting its role as an independent environmental risk factor. Second, when BC exposure occurred together with CRS, depressive-like behavioral alterations became more evident, and selected depression-relevant molecular changes were more prominent in blood and the NAc. This pattern suggests that BC exposure may not only initiate depressive-like behavioral alterations but also worsen stress-related depressive-like outcomes through peripheral and brain-region-specific molecular remodeling. This interpretation is particularly relevant in real-world contexts, where environmental particulate exposure often occurs together with psychosocial stressors rather than in isolation.

Several limitations should be considered. First, this study used a targeted qPCR-based approach, which allowed us to examine selected stress-, inflammation-, and plasticity-related markers but did not provide a genome-wide view of BC-induced molecular alterations. Future transcriptomic or spatially resolved analyses will be useful to define broader molecular networks affected by BC exposure. Second, plasma-depleted whole blood contains mixed blood cell populations; therefore, the observed transcript changes cannot be assigned to specific immune cell types. Future studies using cell-type-specific blood profiling or flow cytometry-based sorting will help identify the cellular sources of these peripheral molecular changes. Third, although we observed coordinated behavioral and molecular alterations, the current design does not establish causal links between specific molecular changes and depressive-like behavior. Genetic or viral manipulation of candidate regulators, such as *Mecp2*, *Bdnf*, or *Creb1*, will be needed to determine whether these molecular changes directly mediate BC-associated behavioral vulnerability. Finally, broader behavioral assessment, protein-level validation, and cell-type-specific brain analyses will be important to determine whether the observed transcript changes translate into functional molecular alterations within defined neural circuits.

In conclusion, our findings provide preclinical evidence that controlled inhaled BC exposure is sufficient to induce depressive-like behavioral alterations and systemic molecular changes. Importantly, BC exposure was associated not only with depressive-like behavioral induction, but also with increased stress-related behavioral and molecular vulnerability when combined with CRS. At the molecular level, BC exposure was linked to peripheral inflammatory and plasticity-related transcript changes, whereas combined BC+CRS exposure revealed selected depression-relevant alterations in blood and the NAc. These results support a model in which BC exposure contributes to depression-related biological vulnerability by reshaping peripheral and brain-region-specific stress response pathways, while concurrent psychological stress further unmasks or intensifies plasticity- and transcription-related molecular changes.

## Acknowledgements

We thank Sun A Jung, Hyunjeong Jin, and Sangjoon Lee for their assistance with the experiments. Some schematic illustrations in this manuscript were created with BioRender.com. This work was supported by the Korea Ministry of Food and Drug Safety (grant no. 25212MFDS003), the National Research Foundation of Korea (grant nos. RS-2024-00332024 and RS-2024-00463082), and KIST (grant no. 26Z9001).

## CRediT authorship contribution statement

**Jinhee Bae:** Conceptualization, Methodology, Investigation, Data curation, Formal analysis, Visualization, Writing – original draft, Writing – review & editing. **Jiwon Lee:** Investigation, Data curation. **Seongeun Song:** Formal analysis, Writing – review & editing. **Kwangil Jeong:** Investigation, Data curation. **Nazarii Frankiv:** Investigation, Data curation. **Chaeran Park:** Investigation, Data curation. **Chae Yeon Hwang:** Investigation, Data curation. **Yun Kyung Kim:** Writing – review & editing. **Byung-Yong Yu:** Conceptualization, Writing – review & editing. **Heh-In Im:** Conceptualization, Supervision, Funding acquisition, Writing – review & editing. All authors reviewed and approved the final manuscript.

## Data availability

The data supporting the findings of this study are included in the article and supplementary information. Additional raw data are available from the corresponding author upon reasonable request.

## Supplementary material

Supplementary material associated with this article can be found in the online version.

