## Supplementary_Material for "Inhaled black carbon induces depressive-like behavior and enhances stress-related blood–brain molecular vulnerability in mice"

Jinhee Bae et al.

This supplementary file contains the following materials:

Supplementary Fig. S1. Additional molecular markers in plasma-depleted whole blood.

Supplementary Fig. S2. Peripheral organ inflammatory transcript profiles following BC exposure and CRS.

Supplementary Fig. S3. Additional molecular markers in stress-related brain regions.

Supplementary Fig. S4. Exploratory analysis of inflammatory transcripts in the mPFC and hippocampus.

Supplementary Table S1. Primer sequences used for quantitative real-time PCR.

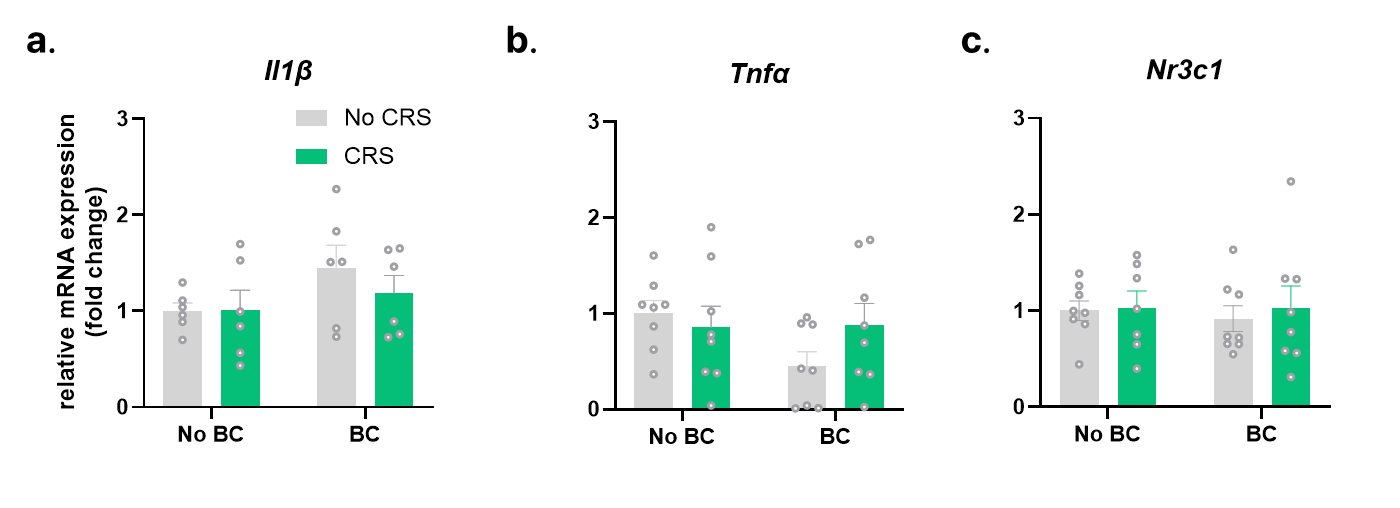
**Supplementary Fig. S1. Additional molecular markers in plasma-depleted whole blood.**
Relative mRNA expression of (a) Il1b, (b) Tnf, and (c) Nr3c1 in plasma-depleted whole blood. No significant changes were detected following BC exposure, CRS, or combined BC+CRS exposure. Data are presented as mean ± SEM. n = 6–8 mice per group.

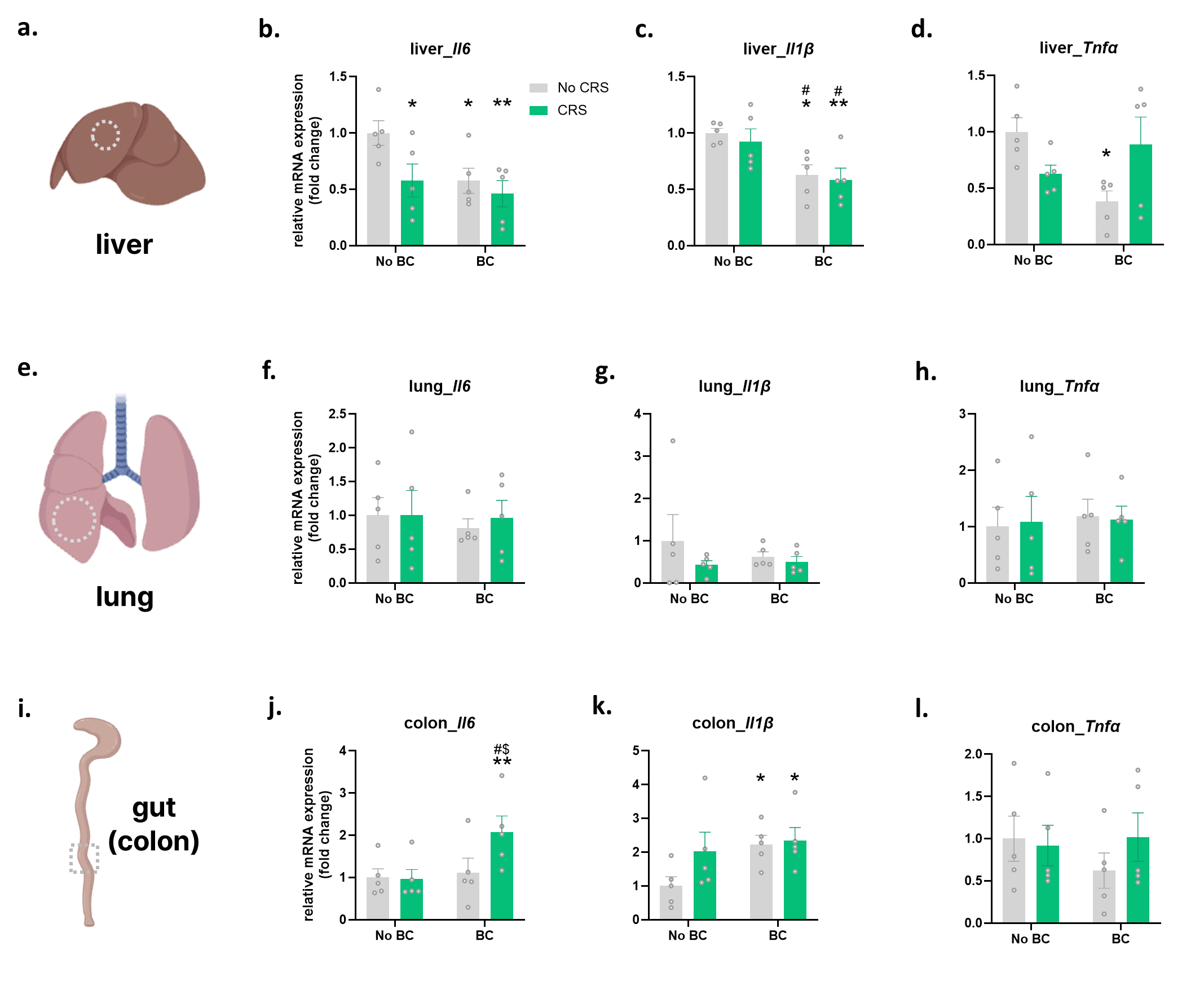

**Supplementary Fig. S2. Peripheral organ inflammatory transcript profiles following BC exposure and CRS.**
(a) Schematic illustration of liver tissue collection. (b–d) Relative mRNA expression of Il6, Il1b, and Tnf in the liver. (e) Schematic illustration of lung tissue collection. (f–h) Relative mRNA expression of Il6, Il1b, and Tnf in the lung. (i) Schematic illustration of colon tissue collection. (j–l) Relative mRNA expression of Il6, Il1b, and Tnf in the colon. Peripheral organ inflammatory transcripts showed organ-specific patterns, with reduced hepatic inflammatory transcript expression, limited lung responses, and elevated colonic Il6 and Il1b expression under BC-exposed or combined exposure conditions. Data are presented as mean ± SEM. n = 5 mice per group. *p < 0.05, **p < 0.01 vs. Control; #p < 0.05 vs. CRS; $p < 0.05 vs. BC.

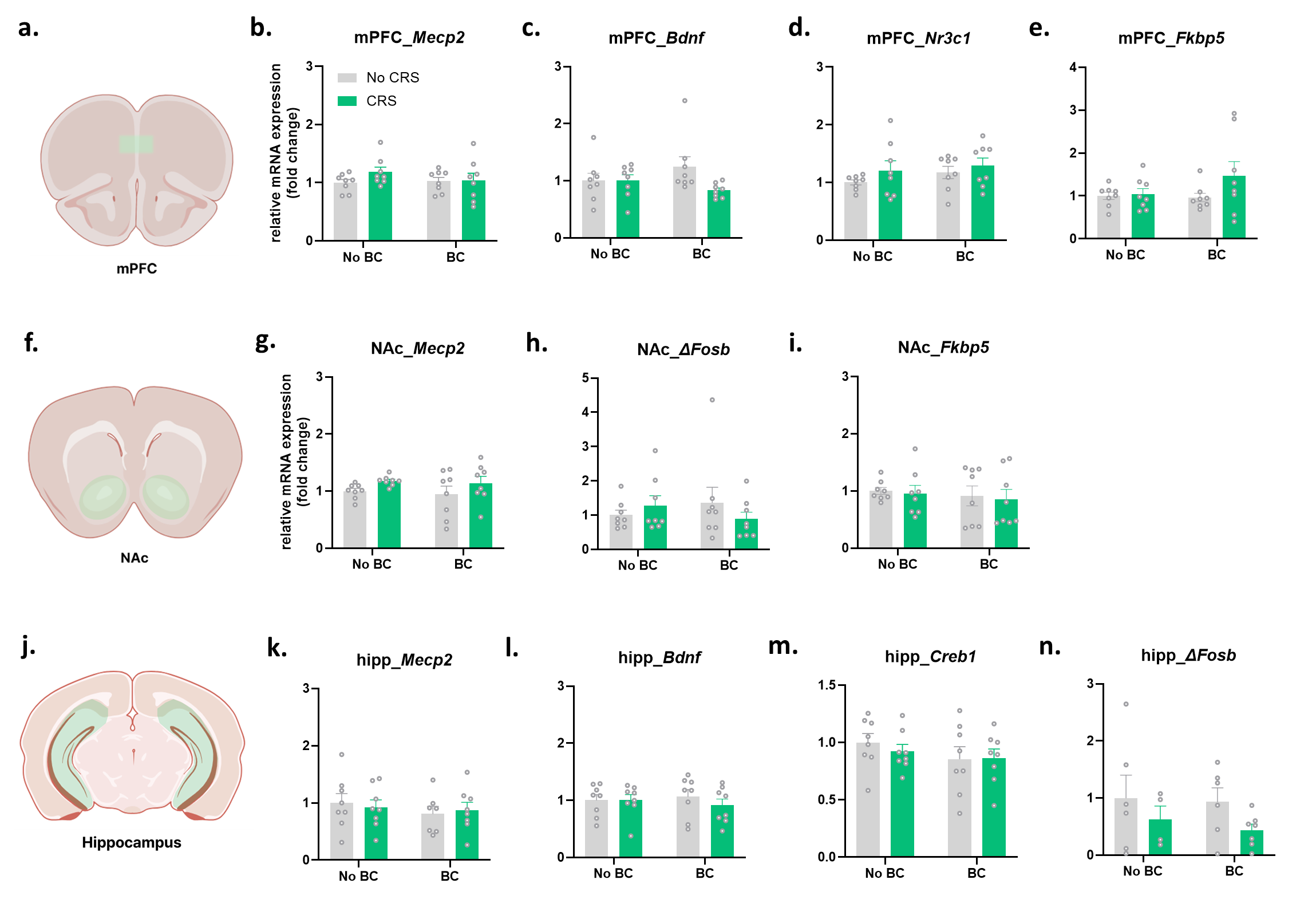

**Supplementary Fig. S3. Additional molecular markers in stress-related brain regions.**
(a) Schematic illustration of the medial prefrontal cortex (mPFC). (b–e) Relative mRNA expression of Mecp2, Bdnf, Nr3c1, and Fkbp5 in the mPFC. (f) Schematic illustration of the nucleus accumbens (NAc). (g–i) Relative mRNA expression of Mecp2, ΔFosB-related Fosb transcript, and Fkbp5 in the NAc. (j) Schematic illustration of the hippocampus. (k–n) Relative mRNA expression of Mecp2, Bdnf, Creb1, and ΔFosB-related Fosb transcript in the hippocampus. These additional screened transcripts did not show robust condition-associated changes across the corresponding brain regions and were included as supplementary molecular data. Data are presented as mean ± SEM. n = 4–8 mice per group.

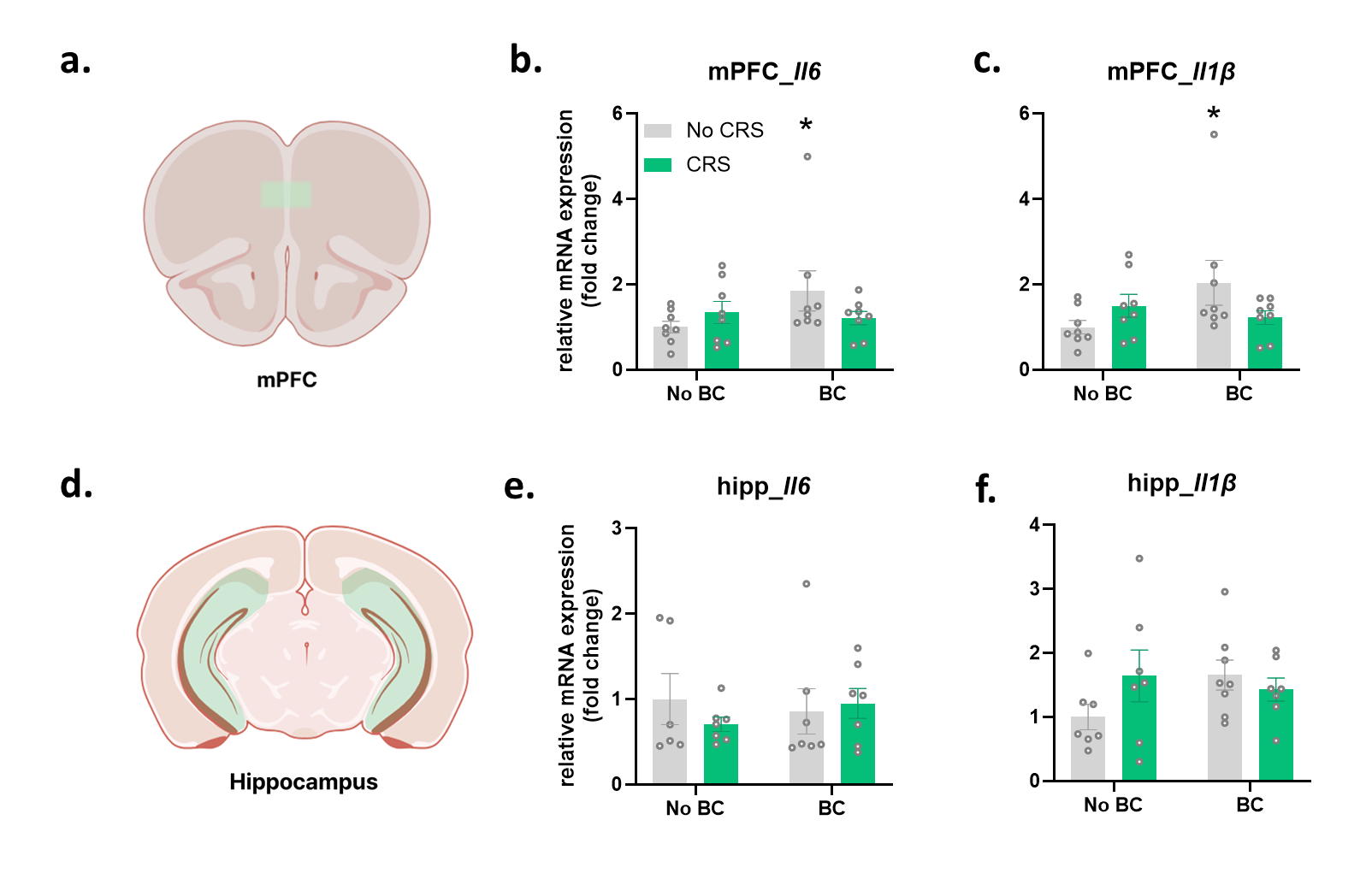

**Supplementary Fig. S4. Exploratory analysis of inflammatory transcripts in the mPFC and hippocampus.**
(a) Schematic illustration of the medial prefrontal cortex (mPFC). (b, c) Relative mRNA expression of Il6 and Il1b in the mPFC. (d) Schematic illustration of the hippocampus. (e, f) Relative mRNA expression of Il6 and Il1b in the hippocampus. Selected mPFC inflammatory transcripts were elevated in the BC-only group, whereas hippocampal inflammatory transcript changes were limited. Data are presented as mean ± SEM. n = 6–8 mice per group. *p < 0.05 vs. Control.

**Supplementary Table S1. Primer sequences used for quantitative real-time PCR.**

| Target transcript | Forward primer sequence (5′–3′) | Reverse primer sequence (5′–3′) |
| --- | --- | --- |
| *Il6* | 5′-TAGTCCTTCCTACCCCAATTTCC-3′ | 5′-TTGGTCCTTAGCCACTCCTTC-3′ |
| *Il1b* | 5′-ACCTAGCTGTCAACGTGTG-3′ | 5′-TCAAAGCAATGTGCTGGTGC-3′ |
| *Tnf* | 5′-ACTGAACTTCGGGGTGATCG-3′ | 5′-GCTTGGTGGTTTGCTACGAC-3′ |
| *Bdnf* | 5′-AGAGCTGTTGGATGAGGACCAG-3′ | 5′-CAAAGGCACTTGACTACTGAGCA-3′ |
| *Creb1* | 5′-CTGAGGAGCTTGTACCACCG-3′ | 5′-CTGCTGGCATGGATACCTGG-3′ |
| *Fkbp5* | 5′-GATTGCCGAGATGTGGTGTTCG-3′ | 5′-GGCTTCTCCAAAACCATAGCGTG-3′ |
| *Nr3c1* | 5′-TGGAGAGGACAACCTGACTTCC-3′ | 5′-ACGGAGGAGAACTCACATCTGG-3′ |
| *Mecp2* | 5′-GAGAGAGCAGAAACCACCTA-3′ | 5′-TCTGATGCTGCTGCCTTT-3′ |
| ΔFosB-related *Fosb* transcript | 5′-ATGTTTCAAGCTTTTCCCGGAGAC-3′ | 5′-CTACTCGGCCAGCGGGC-3′ |
| *Cd3e* | 5′-GCTCCAGGATTTCTCGGAAGTC-3′ | 5′-ATGGCTACTGCTGTCAGGTCCA-3′ |
| *Gapdh* | 5′-GACATCAAGAAGGTGGTGAAGC-3′ | 5′-ACCACCCTGTTGCTGTAGCC-3′ |
